# Structure and heterogeneity of a highly cargo-loaded encapsulin shell

**DOI:** 10.1101/2023.07.26.550694

**Authors:** Seokmu Kwon, Michael P. Andreas, Tobias W. Giessen

**Affiliations:** Department of Chemical Engineering, University of Michigan, Ann Arbor, MI 48109, USA; Department of Biological Chemistry, University of Michigan Medical School, Ann Arbor, MI 48109, USA

**Keywords:** Encapsulin, Nanocompartment, Self-Assembly, Cargo Protein, Structural Heterogeneity

## Abstract

Encapsulins are self-assembling protein nanocompartments able to selectively encapsulate dedicated cargo enzymes. Encapsulins are widespread across bacterial and archaeal phyla and are involved in oxidative stress resistance, iron storage, and sulfur metabolism. Encapsulin shells exhibit icosahedral geometry and consist of 60, 180, or 240 identical protein subunits. Cargo encapsulation is mediated by the specific interaction of targeting peptides or domains, found in all cargo proteins, with the interior surface of the encapsulin shell during shell self-assembly. Here, we report the 2.53 Å cryo-EM structure of a heterologously produced and highly cargo-loaded T3 encapsulin shell from *Myxococcus xanthus* and explore the systems’ structural heterogeneity. We find that exceedingly high cargo loading results in the formation of substantial amounts of distorted and aberrant shells, likely caused by a combination of unfavorable steric clashes of cargo proteins and shell conformational changes. Based on our cryo-EM structure, we determine and analyze the targeting peptide-shell binding mode. We find that both ionic and hydrophobic interactions mediate targeting peptide binding. Our results will guide future attempts at rationally engineering encapsulins for biomedical and biotechnological applications.

## Introduction

Encapsulins are cargo-loaded protein nanocompartments found in a wide variety of prokaryotes (Giessen, 2022). Encapsulin systems have been implicated in oxidative stress resistance (Lien et al., 2021; Tang et al., 2021; Tracey et al., 2019), sulfur metabolism (Benisch et al., 2023; Nichols et al., 2021), iron storage (Eren et al., 2022; Giessen et al., 2019; He et al., 2016; LaFrance et al., 2021; Piergentili et al., 2020), and the biosynthesis of secondary metabolites (Andreas and Giessen, 2021; Andreas and Giessen, 2022). Encapsulin shells self-assemble – even in the absence of cargo – into icosahedral protein compartments with diameters between 18 and 42 nm and varying triangulation numbers (T1: 60 subunits, T3: 180 subunits, or T4: 240 subunits) (Giessen, 2022). Encapsulin shell proteins possess the HK97 phage-like fold and may have originated from defective prophages whose capsid components have been coopted by the ancestral cellular host (Andreas and Giessen, 2021; Giessen and Silver, 2017; Nichols et al., 2017). Encapsulins can be grouped into four distinct families based on sequence similarity, operon organization, domain architecture, and encapsulation mechanism (Andreas and Giessen, 2021). Family 1 is the most extensively characterized while Family 2 has only recently begun to be studied experimentally. Family 3 and Family 4 remain putative and currently lack experimental validation. The eponymous feature of encapsulins is their ability to encapsulate specific cargo proteins during shell self-assembly (Altenburg et al., 2021; Cassidy-Amstutz et al., 2016; Sutter et al., 2008; Tamura et al., 2015). Cargo sequestration inside the shell is mediated by selective cargo loading mechanisms based on targeting domains (TDs) or targeting peptides (TPs) present at the N- or C-terminus of all cargo proteins (Jones et al., 2023a). This native, efficient, and modular cargo loading modality makes encapsulins excellent engineering platforms for biomedical and biotechnological applications ranging from targeted drug delivery and vaccine platforms to bionanoreactors and novel biomaterials (Gabashvili et al., 2022; Jenkins and Lutz, 2021; Jones and Giessen, 2021; Jones et al., 2021; Lagoutte et al., 2018; Van de Steen et al., 2021).

While little information regarding TDs and their cargo loading mechanism is currently available, TP-based cargo loading is better understood. The TP binding site has been determined to reside on the inner surface of the encapsulin shell protein (Jones et al., 2023a). The maximal number of cargo proteins that can be specifically loaded into a given encapsulin is determined by the total number of shell protein subunits, also called protomers, each providing one binding site – 60 for T1, 180 for T3, and 240 for T4 shells. In native and heterologous encapsulins, the actual amount of cargo loaded is likely dependent on a combination of factors including triangulation number, TP binding strength, cargo size and oligomerization state, the kinetics of shell assembly, and the relative expression levels of shell and cargo proteins (Altenburg et al., 2021). TPs in native cargo proteins are usually separated from the globular catalytic cargo domain by a 10-50 residue long flexible linker which likely helps minimize steric clashes between cargo proteins bound to adjacent TP binding sites (Giessen, 2022). A corollary of this feature is that cargo proteins are usually poorly resolved in structures of cargo-loaded encapsulin shells due to their high mobility (McDowell and Hoiczyk, 2022). Low cargo loading and correspondingly low TP occupancy or the simultaneous loading of multiple different cargos with distinct TPs, generally results in weak or hard to interpret TP densities in cryo-EM maps, making it difficult to build high-confidence atomic models to explore the details of TP-shell interactions. Specifically, in the Family 1 model T3 encapsulin from *Myxococcus xanthus* (Mx) (McHugh et al., 2014), the natively observed mixed loading of three distinct cargo proteins and the low cargo loading observed for individual cargos in heterologous expression experiments, have so far only yielded TP densities of moderate quality, leading to difficulties in building high-confidence atomic models (Eren et al., 2022).

A given cargo-loaded encapsulin usually assembles into only one defined shell size with either T1, T3, or T4 icosahedral geometry (Giessen, 2022). In the absence of cargo, presence of non-native cargo, or artificially induced high cargo-loading conditions, the Mx encapsulin has been observed to form mixtures of T1 and T3 shells (Eren et al., 2022; McHugh et al., 2014). Even though this structural polymorphism is established in the literature, little is known about the underlying mechanistic details. Further, a small percentage of aberrant assemblies can often be observed under the non-native assembly conditions listed above. So far, little attention has been paid to these assemblies.

In this study, we show that heterologous loading of a small monomeric non-native cargo protein into the Mx encapsulin shell results in high cargo occupancy (ca. 95%). We further show that high cargo loading leads to increased structural heterogeneity of the protein shell. Additionally, we report the 2.53 Å cryo-EM structure of the highly cargo-loaded Mx T3 shell with improved targeting peptide (TP) densities, allowing confident atomic model building and revealing the details of the TP-shell interactions in the Mx T3 model encapsulin.

## Methods

### Molecular cloning

DNA constructs of Mx EncA and SNAP-tag-TP were ordered from Integrated DNA Technologies (IDT) as *E. coli* codon-optimized gBlock fragments (Table S1). The expression plasmid was constructed through successive Gibson assembly of the two genes using linearized pET-Duet1 vector obtained via PCR, resulting in an expression plasmid encoding Mx EncA in MCS1 and SNAP-tag-TP in MCS2. *E. coli* BL21 (DE3) cells were transformed with the assembled plasmid via electroporation and the expression strain confirmed through Sanger sequencing (Eurofins Scientific).

### Protein production and purification

Protein production and purification was carried out as previously described (Kwon and Giessen, 2022). Briefly, all expression experiments were carried out using lysogeny broth (LB) medium supplemented with 100 mg/mL ampicillin. 500 mL of fresh LB medium were inoculated 1:100 using a 5 mL overnight culture, grown at 37°C to an OD_600_ of 0.5-0.6, and induced with 0.2 mM IPTG. After induction, cultures were grown at 30°C overnight for 18 h and harvested via centrifugation (5,000 g, 12 min, 4°C) and stored at -20°C for later use. Frozen cell pellets were resuspended in 5 mL/g (wet cell mass) of Tris buffer (20 mM Tris, 150 mM NaCl, pH 7.5). Lysozyme (0.5 mg/mL), Benzonase nuclease (25 units/mL), and MgCl_2_ (1.5 mM) were added followed by sonication (Model 120 Sonic Dismembrator, Fisher Scientific) at 60% amplitude for 5 min total (10 seconds on, 20 seconds off) until no longer viscous. After sonication, samples were clarified by centrifugation (10,000 g, 15 min, 4°C) and the supernatant subjected to polyethylene glycol (PEG) precipitation. After centrifugation (5,000 g, 12 min, 4°C), the resulting PEG pellet was resuspended in 3 mL Tris buffer (pH 7.5) and filtered using a 0.2 mm syringe filter. The filtered sample was subjected to size exclusion chromatography (SEC) using a Sephacryl S-500 16/60 column and Mx EncA-containing fractions were combined, concentrated, and dialyzed using Amicon filter units (100 kDa) and Tris buffer (20 mM Tris, pH 7.5). The low salt sample was then loaded on a HiPrep DEAE FF 16/10 Ion Exchange column for ion exchange chromatography. Mx EncA-containing fractions were concentrated, centrifuged (10,000 g, 10 minutes, 4°C), and subjected to a second round of SEC using a Superose 6 10/300 GL column and Tris buffer (20 mM Tris, 150 mM NaCl, pH 7.5). Protein fractions were again collected and purified protein was stored in Tris buffer (20 mM Tris, 150 mM NaCl, pH 7.5) at 4°C until further use.

### SDS polyacrylamide gel electrophoresis (SDS-PAGE)

SDS polyacrylamide gel electrophoresis was conducted in an Invitrogen XCell SureLock using Novex 14% Tris-Glycine Mini Protein Gels with SDS running buffer. 4.6 μg of each sample was mixed with 4X SDS sample buffer, heated for 5 min at 98°C, centrifuged (10,000 g, 2 min), and then loaded onto the gel. SDS PAGE analysis was carried out at a constant voltage of 225 V for 42 min at room temperature. The Spectra Multicolor Broad Range Protein Ladder was used.

### Native polyacrylamide gel electrophoresis (native PAGE)

Native polyacrylamide gel electrophoresis was conducted in an Invitrogen XCell SureLock using NativePAGE 3 to 12% bis-tris mini protein gels with 1X NativePAGE Anode Buffer and 1X NativePAGE Cathode Buffer. 5.2 μg of each sample was mixed with 4X NativePAGE sample buffer and then loaded onto the gel. Native PAGE was carried out at a constant voltage of 150 V for 1 hour followed by an additional 1 hour run at 250 V at 4°C. NativeMark Unstained Protein Standard was used as a protein ladder. After the run, gel was soaked in water to remove dye, and then stained with Coomassie Blue for protein detection.

### Negative stain transmission electron microscopy (TEM)

Encapsulin samples for negative-stain TEM were diluted to 0.15 mg/mL in Tris buffer (20 mM Tris, 150 mM NaCl, pH 7.5). Gold TEM grids (200-mesh coated with Formvar-carbon film, EMS) were glow discharged at 5 mA for 60 s (easiGlow, PELCO). 4 mL of sample was added to the grid and incubated for 1 min, excess liquid wicked with filter paper, and washed with 0.75% uranyl formate before staining with 0.75% uranyl formate for 1 min. Stain was removed using filter paper and the grid was dried for at least 20 min before imaging. TEM micrographs were captured using a Morgagni transmission electron microscope at 100 keV.

### Cryo-EM sample preparation and data collection

Grids were prepared by applying 3.5 mL of freshly prepared protein at a concentration of 3.4 mg/mL to glow-discharged Quantifoil R2/1, 200 mesh copper holey carbon grids (EMS, CAT: Q225CR1). The grids were then frozen by plunging into liquid ethane using an FEI Vitrobot Mark IV with the following parameters: temperature 22°C, humidity 100%, blot force 5, blot time 2 seconds. The grids were clipped and stored in liquid nitrogen until data collection.

### Cryo-EM data collection

Cryo-EM movies were collected using an FEI Titan Krios G3 cryo-transmission electron microscope operating at 300 kV and equipped with a Gatan K3 Direct Detector with Bioquantum Imaging Filter. SerialEM (Mastronarde, 2003) was used to select targets and acquire movies with the following settings: defocus range -1 to -2.5 μm, dose 49.3 e^-^/Å^2^, magnification 105,000x, exposure time 3.2 seconds with 63.9 milliseconds per frame. 2,610 movies were collected in total from a single grid.

### Cryo-EM data processing

All data processing was carried out using CryoSPARC v4.2.1 (Punjani et al., 2017) (Figure S1). 2,610 movies were motion-corrected using patch motion correction, followed by patch CTF estimation. Movies with CTF fits worse than 5 Å were discarded, resulting in 2,440 accepted movies. Template picker was used for particle picking. 96,383 particles were extracted with a box size of 560 pixels. The particles were then downsampled to a box size of 440 pixels and sorted with two rounds of 2D classification, picking only classes containing well-resolved T3 shells which resulted in 63,680 particles. The particles were then used to generate two ab-initio volumes with I symmetry imposed, with the majority volume containing 63,543 particles. The particles from the majority ab-initio volume were used in an non-uniform (NU) refinement with I symmetry, per-particle defocus optimization, per-group CTF parameter optimization, and Ewald sphere correction applied, resulting in volume with a global resolution of 2.57 Å (Punjani et al., 2020). Then, heterogenous refinement with three classes was performed using three copies of the NU-refined map as initial volumes, yielding a majority class containing 50,680 particles. These particles were used in another NU refinement with I symmetry, per-particle defocus optimization, per-group CTF parameter optimization, and Ewald sphere correction applied to produce a final map with 2.53 Å resolution. Local resolution estimation was carried out using CryoSPARC.

### Atomic model building, refinement, and structural analysis

To build the Mx T3 shell, a previously published model of the T3 Mx encapsulin ASU with targeting peptides (PDB ID: 7S2T) was docked into the cryo-EM map using ChimeraX v 1.15 (Pettersen et al., 2021) by using the fit-to-volume command. The placed coordinates were then manually refined against the map using Coot v 0.9.8.1 (Emsley et al., 2010) until the fit was satisfactory. Phenix v 1.20.1-4487 (Liebschner et al., 2019) was used to find the symmetry operators from the map using the symmetry-from-map function, which were then used to generate a T3 icosahedral shell containing 180 copies of the Mx EncA protomer with 180 targeting peptides. The model was refined using real-space refinement with NCS restraints, three macro cycles, and all other settings left to default. The model was further improved by iterative manual refinements using Coot. The final model was validated against the map using the comprehensive validation tool in Phenix to ensure that the model’s geometry and map fit were satisfactory. The model was deposited in the Protein Data Bank (PDB) under PDB ID: 8TK7; and the Electron Microscopy Data Bank (EMDB) under EMD-41322.

## Results

### Heterologous cargo loading results in high cargo occupancy

In this study, we focus on the model T3 encapsulin system from *M. xanthus* DK 1622 which is natively expressed at low levels during vegetative growth and strongly induced under starvation (McHugh et al., 2014). The Mx T3 encapsulin has been shown to protect *M. xanthus* from oxidative stress through iron sequestration and to bind, condense, and stabilize plasmid DNA *in vitro* (Almeida et al., 2022). Further, mutants of the Mx T3 shell protein (EncA) are likely tan-phase-locked, defective in the production of the secondary metabolites DKxanthene and myxovirescin, agglutination of cells, extracellular polysaccharide production, and predation, indicating that EncA plays an important role in the developmental life cycle of *M. xanthus* (Kim et al., 2019). EncA natively assembles into a T3 shell loaded with three different cargo proteins (EncB, EncC, and EncD), two of them – EncB and EncC – belonging to the ferritin-like protein (Flp) superfamily (Fig. 1a) (McHugh et al., 2014). EncB and EncC have been shown to assemble into five-fold symmetrical decameric ring-shaped complexes consisting of ten two-helix bundle monomers each (Eren et al., 2022). EncD has not been characterized so far and shows no homology to characterized proteins. The relative loading observed in natively isolated Mx encapsulins is 52% EncC, 27% EncD, and 21% EncB (Fig. 1a).

**Fig. 1.**
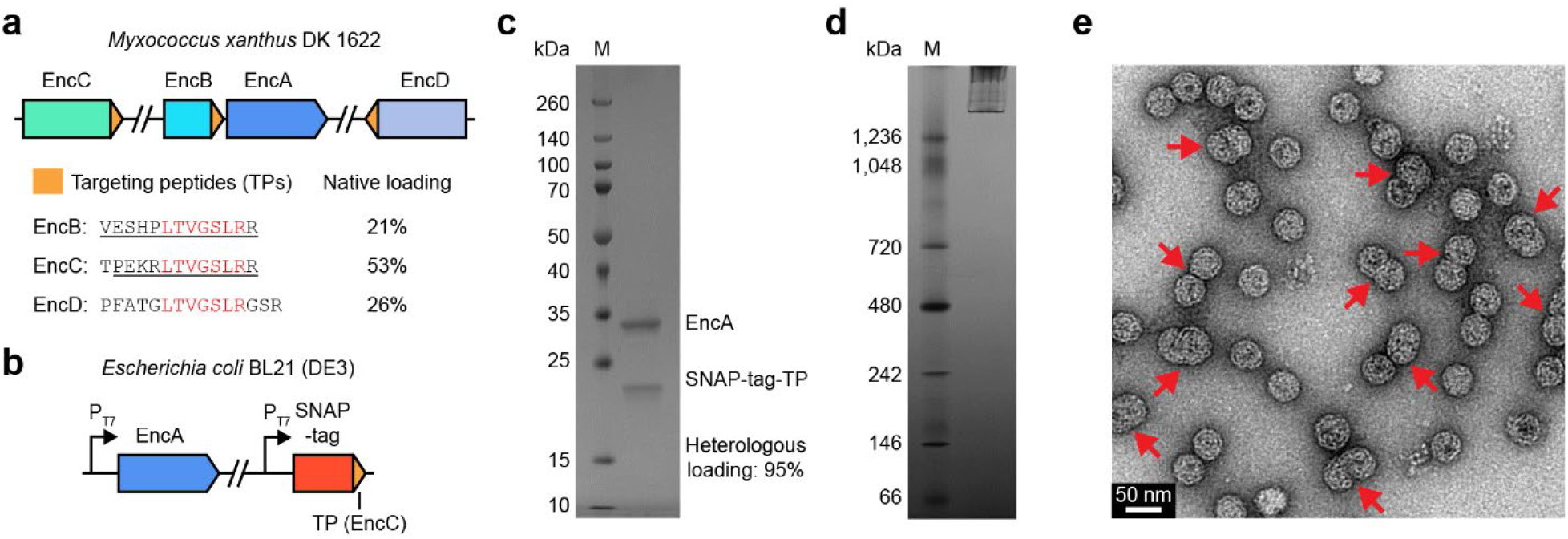
Heterologous production of highly cargo-loaded encapsulin shells. (**a**) The *Myxococcus xanthus* Family 1 encapsulin operon (top) and targeting peptides (TPs) of the three known cargo proteins EncB, EncC, and EncD (bottom). Conserved TP residues are highlighted in red. Underlined residues were previously shown to be involved in cargo loading. The native cargo loading percentages observed in encapsulins purified from *M. xanthus* are shown (McHugh et al., 2014). (**b**) Heterologous expression strategy for producing highly cargo-loaded Mx encapsulins. EncA and the cargo protein SNAP-tag-TP are under the control of separate T7 promoters. (**c**) SDS-PAGE gel of purified cargo-loaded EncA shells. Heterologous cargo loading based on gel densitometry is shown. (**d**) Native PAGE gel of purified SNAP-tag-TP-loaded EncA shells highlighting the absence of any lower molecular weight bands, thus confirming cargo encapsulation. (**e**) Representative negative stain TEM micrograph of purified cargo-loaded EncA shells. Distorted and fusion shells deviating from the expected T3 size (32 nm) and geometry are highlighted by red arrows.

This heterogeneous cargo loading mode combined with the initially achieved moderate resolution resulted in difficulties in determining the details of TP-shell interaction via cryo-EM. Using EncB and EncC individually in heterologous cargo loading experiments resulted in ca. 40-50% cargo loading which allowed the visualization of bound TPs (Eren et al., 2022). The observed densities exhibited improved but still limited resolution. To achieve high and homogeneous cargo loading – and correspondingly strong and high resolution TP densities – we heterologously expressed the Mx encapsulin shell protein EncA together with a small, monomeric, non-native cargo protein (SNAP-tag-TP) in *E. coli* BL21 (DE3). The TP found in the native main cargo EncC (PEKRLTVGSLRR) was fused to the C-terminus of SNAP-tag, separated from the globular cargo domain by an eight residue linker (GGGSGGGS) to prevent steric clashes between cargo bound to neighboring TP binding sites (Table S1). Our expression strategy was based on a single plasmid (pETDuet-1) encoding the shell and cargo proteins in different loci, each under the separate control of a strong T7 promoter and a strong *E. coli* ribosome binding site (Fig. 1b). Heterologous expression, followed by purification via polyethylene glycol precipitation, gel filtration on a Sephacryl S-500 column, ion exchange chromatography, and gel filtration on a Superose 6 column, yielded cargo-loaded and readily assembled encapsulin shells as confirmed by SDS-PAGE, native PAGE, and negative stain transmission electron microscopy (TEM) (Fig. 1c-e). Cargo loading was calculated based on gel densitometry taking the relative molecular weights of EncA (31.7 kDa) and SNAP-tag-TP (21.4 kDa) into account and was determined to be ca. 95% (Fig. 1c). Negative stain TEM micrographs showed a substantial number of aberrant non-T3 shells (Fig. 1e, red arrows), with some appearing to be fusions of two T3 shells while others appeared to be expanded in size or distorted.

### Structural polymorphism and heterogeneity in highly loaded encapsulins

To further investigate the structure and apparent heterogeneity of the highly cargo-loaded Mx encapsulin shell, we carried out cryo-EM experiments. Cryo-EM micrographs confirmed that the aberrant structures observed in negative stain TEM were not sample preparation or imaging artifacts as they were clearly visible in vitrified samples as well (Fig. 2a, arrows). Similar to negative stain micrographs, cryo-EM images also contained a substantial number of fused, elongated, or expanded shells, in addition to normal T3 shells. We define fusion shells as two-lobed particles with a long axis of 48 nm or longer (Fig. 2a, red arrows) – 150% of the T3 shell diameter of 32 nm. 2D classification of fusion particles did not yield high-resolution classes, likely due to their size and shape heterogeneity. 2D classification of non-fusion particles resulted in a number of well-resolved classes representing T3 shells and distorted shells that do not exhibit clear T3 or T4 icosahedral geometry (Fig. 2b, orange arrows). Approximately 20% of non-fusion particles could be sorted into 2D classes exhibiting distorted shells. Asymmetric (C1) ab-initio reconstructions of a subset of distorted 2D classes yielded irregular closed shells containing apparent pentameric and hexameric facets, without the regular tiling observed in icosahedral T3 or T4 shells (Fig. 2c) (Giessen, 2022).

**Fig. 2.**
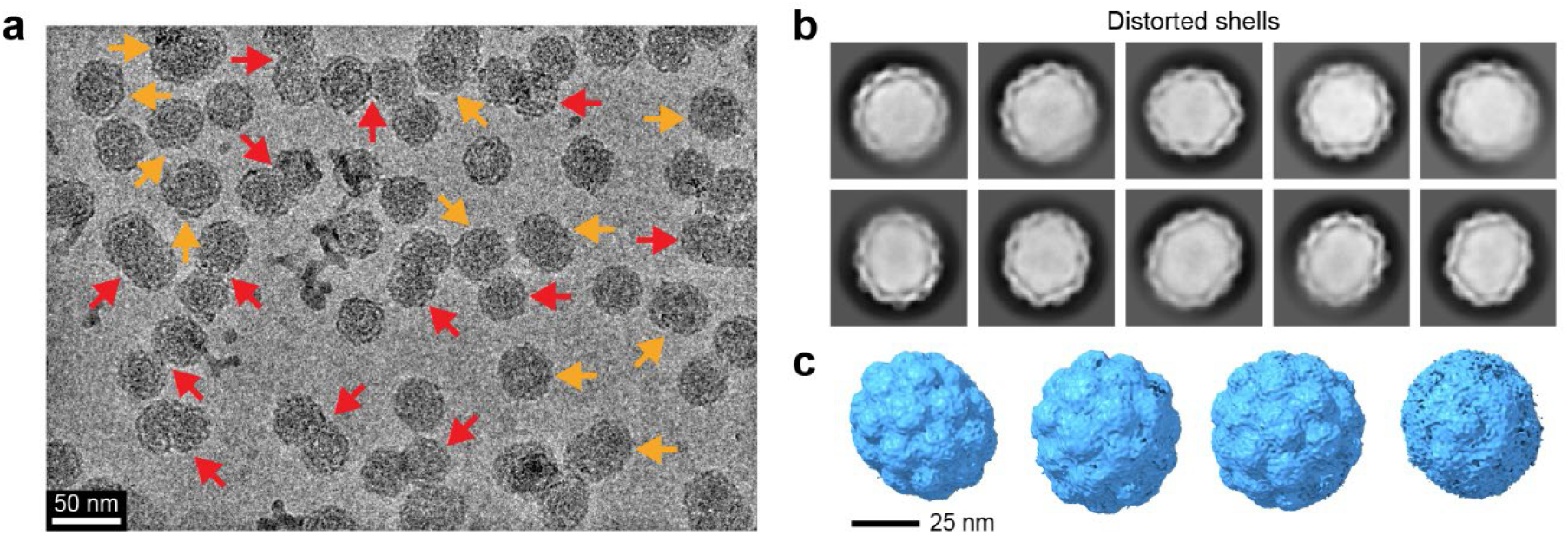
Structural polymorphism and heterogeneity of highly cargo-loaded encapsulin shells. (**a**) Representative cryo-EM micrograph of SNAP-tag-TP-loaded EncA shells. Fusion shells (red arrows) and distorted shells (orange arrows) are highlighted. Fusion shells are defined as two-lobed particles with a long axis of 48 nm or longer while distorted shells encompass all other aberrant shells deviating from icosahedral T3 shells. (**b**) Representative 2D class averages of distorted Mx shells loaded with SNAP-tag-TP. (**c**) Select C1 ab-initio volumes based on particles contained in the 2D classes shown in **b**.

### Single particle cryo-EM analysis of a highly cargo-loaded T3 shell

To gain molecular level insight into the structure of the highly cargo-loaded Mx T3 shell and TP-shell interactions, single particle cryo-EM analysis was carried out (Figure S1). The T3 encapsulin shell was determined to 2.53 Å via icosahedral (I) refinement (Fig. 3a,b). As suggested by negative stain TEM, the T3 encapsulin shell was found to be 32 nm in diameter and to contain

**Fig. 3.**
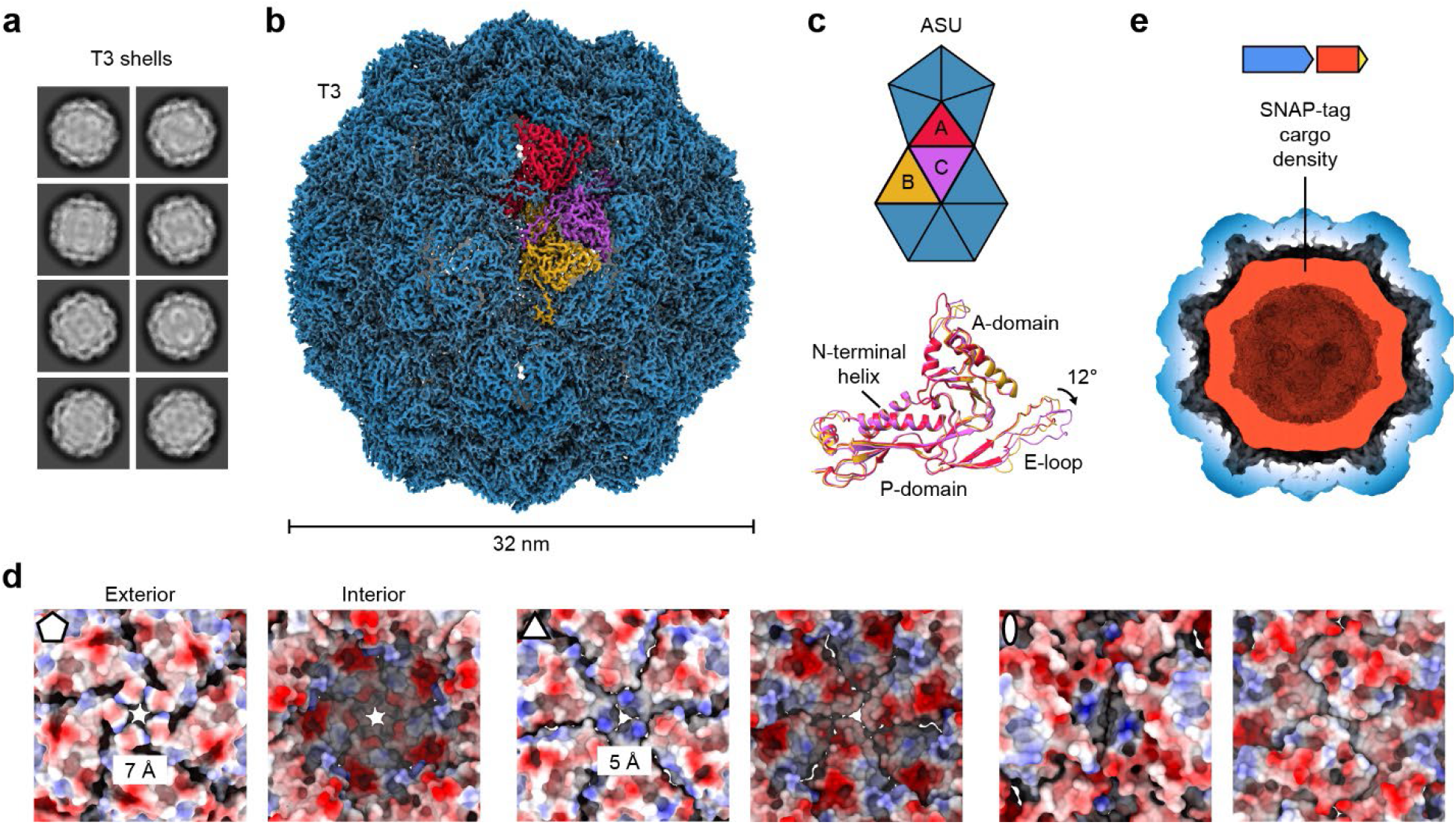
Single particle cryo-EM analysis of highly cargo-loaded T3 encapsulins. (**a**) Representative 2D class averages of the SNAP-tag-TP-loaded EncA T3 shell. (**b**) Cryo-EM map of the Mx T3 shell. The asymmetric unit (ASU) of the T3 shell is highlighted in color. (**c**) ASU diagram (top) and structural alignment of the three unique protomers (A, B, and C) found in the ASU (bottom). The observed 12° shift of the E-loop of the C protomer relative to protomers A and B is highlighted. (**d**) Exterior and interior views of the 5-, 3-, and 2-fold pores of the T3 shell shown in electrostatic surface representation. The pore diameters of the 5- and 3-fold pores are shown. (**e**) Gaussian blurred (ChimeraX: sdev 2) cryo-EM map of the SNAP-tag-TP-loaded T3 shell highlighting internal cargo density (Pettersen et al., 2021).

180 protomers, consistent with previously reported T3 encapsulin structures (Akita et al., 2007; Eren et al., 2022; McHugh et al., 2014). The T3 asymmetric unit (ASU) contains three protomers, one part of a pentameric vertex (protomer A) and two part of a hexameric facet (protomers B and C) which exhibit the canonical HK97 phage-like fold consisting of an A-domain (axial domain), P-domain (peripheral domain), and E-loop (extended loop) (Fig. 3b,c) (Duda and Teschke, 2019). Structural alignments of the three unique ASU protomers indicate that the two primary locations of structural difference are the E-loop and the loop at the tip of the A-domain. Protomers A and B exhibit nearly identical E-loop conformations while differing in the orientation of the A-domain loop. Protomers B and C show similar A-domain loops while the E-loop of C is tilted forward by ∼12° compared to the E-loops of A and B (Fig. 3c). The overall RMSD values between unique protomers are 2.1 Å (A,B), 2.6 Å (A,C), and 2.0 Å (B,C).

Encapsulin shells contain pores at the 5-, 3-, and 2-fold axes of symmetry. Different pore sizes have been observed in different encapsulins ranging from <1 to 20 Å (Giessen, 2022; Jones et al., 2023b; Wiryaman and Toor, 2022). In addition, the 5-fold pores of some encapsulins were found to be dynamic, existing in distinct open and closed states (Ross et al., 2022). Pores are thought to be essential for allowing the substrates, cofactors, and products of encapsulated cargo enzymes to selectively and efficiently cross the shell (Adamson et al., 2022). In the Mx T3 shell, pores are likely necessary for allowing the flux of ferrous iron – the substrate of the encapsulated Flp cargo enzymes EncB and EncC – into the encapsulin (Eren et al., 2022). We observed two main types of open pores in the Mx T3 shell located at the 5- and 3-fold axes of symmetry, positioned at the center of all shell pentamers (7 Å) and hexamers (5 Å). In contrast, the 2-fold pores were found to be completely closed (Fig. 3d). In total, there are 32 open pores in the Mx T3 shell. Based on surface electrostatics, the 5-fold pore exhibits negatively charged and neutral surface patches on its exterior and interior, respectively, and is relatively neutral at its narrowest point. The 3-fold pore is positively charged on its exterior and narrowest point and negatively charged on the interior. Based on pore charge, the 5-fold pore has previously been suggested to represent the entry point to the shell lumen for the ferrous iron substrate of the Flp cargo enzymes (Eren et al., 2022). This model is in good agreement with the observed pore charges in our Mx T3 shell structure.

Cryo-EM density of the SNAP-tag-TP cargo can be visualized inside the Mx T3 shell (Fig. 3e). The cryo-EM map of the cargo-loaded Mx T3 encapsulin clearly exhibits substantial symmetry-averaged internal density – not present in empty shells (Eren et al., 2022). Similar low resolution density was found in asymmetric C1 refinements (data not shown). This density corresponds to the SNAP-tag-TP cargo proteins and appears disconnected from the shell. SNAP-tag-TP is flexibly tethered to the shell interior via direct interaction between the TP and its binding site. However, the globular SNAP-tag domain itself is highly mobile due to the flexible and unresolvable linker connecting it to the TP, preventing high-resolution visualization of cargo density and resulting in cargo density appearing disconnected from the shell.

### Structural details of cargo encapsulation in a T3 encapsulin

The cryo-EM map of the highly cargo-loaded Mx T3 shell exhibits strong high quality TP density (Fig. 4a). TP density can be found in both pentameric and hexameric facets. Local resolution analysis highlights that all TPs are well resolved with resolutions similar to the shell at around 2.5 Å (Fig. 4b).

**Fig. 4.**
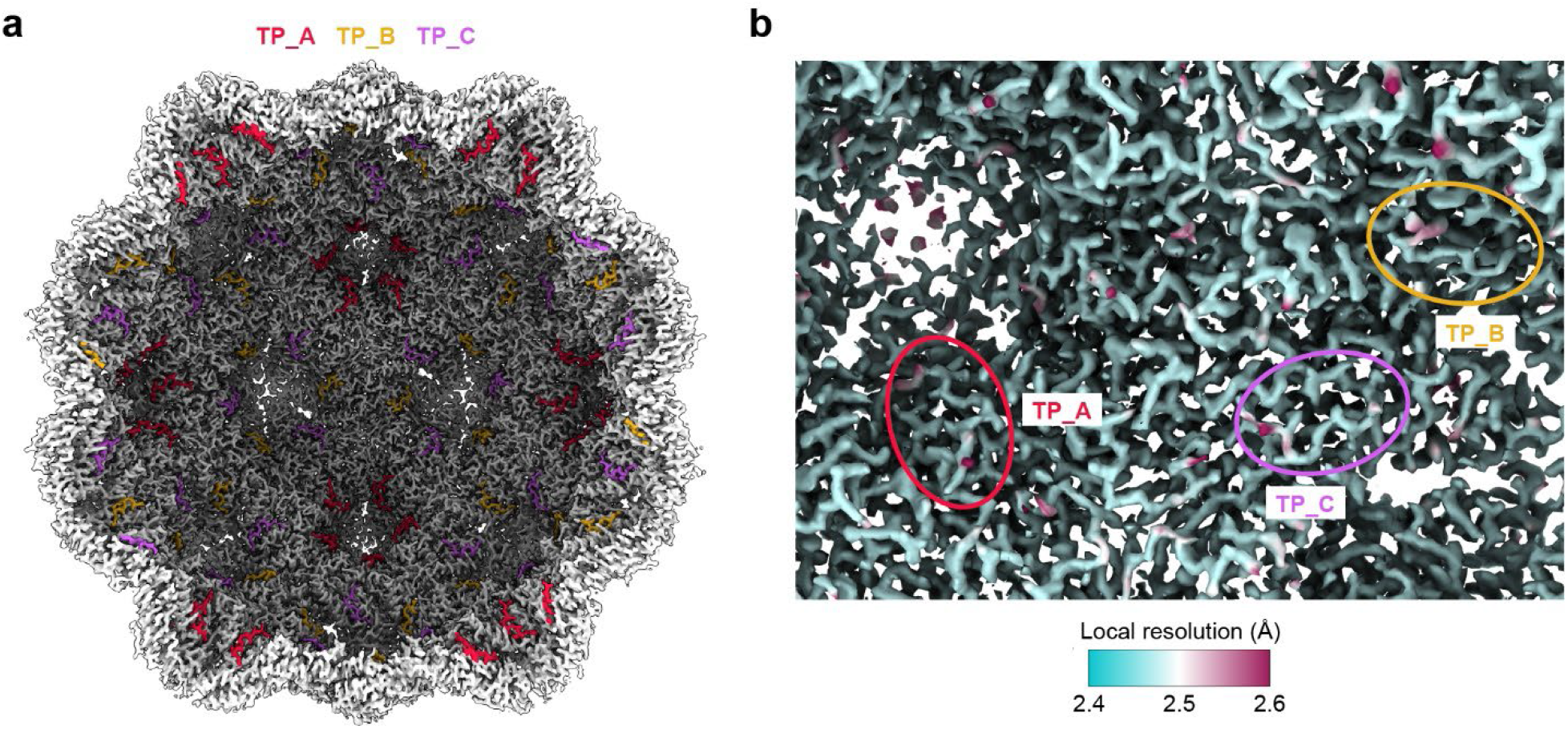
Structural basis for cargo loading in T3 encapsulins. (**a**) Cut-away view of the T3 encapsulin showing the shell interior. The three distinct TPs are highlighted in color (TP_A binding to protomer A of the ASU, TP_B to protomer B, and TP_C to protomer C). (**b**) View onto the interior surface of the T3 shell. The cryo-EM map is colored by local resolution. The three unique TPs and their binding sites are highlighted.

This allowed for high-confidence atomic model building of all three unique TPs (TP_A, TP_B, and TP_C) occupying the binding sites of the ASU protomers A, B, and C. Ten TP residues – EKRLTVGSLR – could be confidently modeled into all three unique TP densities (Fig. 5a). This is in agreement with previous studies and bioinformatic analyses that have shown that TPs in Family 1 encapsulins usually contain one or two conserved GSL motifs, often followed by one or two positively charged residues (Andreas and Giessen, 2021; Cassidy-Amstutz et al., 2016; Jones et al., 2023a). Structural alignments of the three unique TPs show that their conformations are highly similar with the C-terminal ends being nearly identical while the N-terminal ends appear to be more flexible (Fig. 5b,c). Overall, the three TPs possess RMSD values of 1.7 Å (A,B), 1.0 Å (A,C), and 2.0 Å (B,C). As observed for other encapsulin systems, the TP binding site is formed by the P-domain and N-terminal helix and is located on the interior of each shell protein for a total of 180 TP binding sites per Mx T3 shell (Fig. 4a). Recently, a general TP binding mode has been proposed where two or three hydrophobic residues, spaced one or two residues apart, interact with hydrophobic patches within the binding pocket while the C-terminal positively charged residue(s) interact with negatively charged surface patches (Jones et al., 2023a). Our structure supports and expands this binding mode (Fig. 5d,e). As previous models of TP-shell interaction (Eren et al., 2022) were based on lower resolution and less defined cryo-EM densities, specific ionic and hydrogen bonding could not be assigned (Fig. 5d). Our structure highlights a number of likely ionic and H-bonding interactions between TP residues (EKRL**TVGS**L**R**, interacting residues in bold) and shell protomer side chains (R28, R34, D38, D213, D243). In addition, an interaction between a TP lysine residue (E**K**RLTVGSLR) and a backbone carbonyl (N105) of a neighboring shell protomer could be observed. Three hydrophobic residues (EKR**L**_**(1)**_T**V**GS**L**_**(2)**_R, hydrophobic residues in bold) interact with the hydrophobic pockets P1 (**L**_**(1)**_), P2 (**V**), and P3 (**L**_**(2)**_) (Fig. 5e). P1 is located opposite to P2 and P3, an unusual arrangement as all other previously determined TP-shell interactions rely on two or three hydrophobic pockets localized on one – the P2/P3 – side (Jones et al., 2023a).

**Fig. 5.**
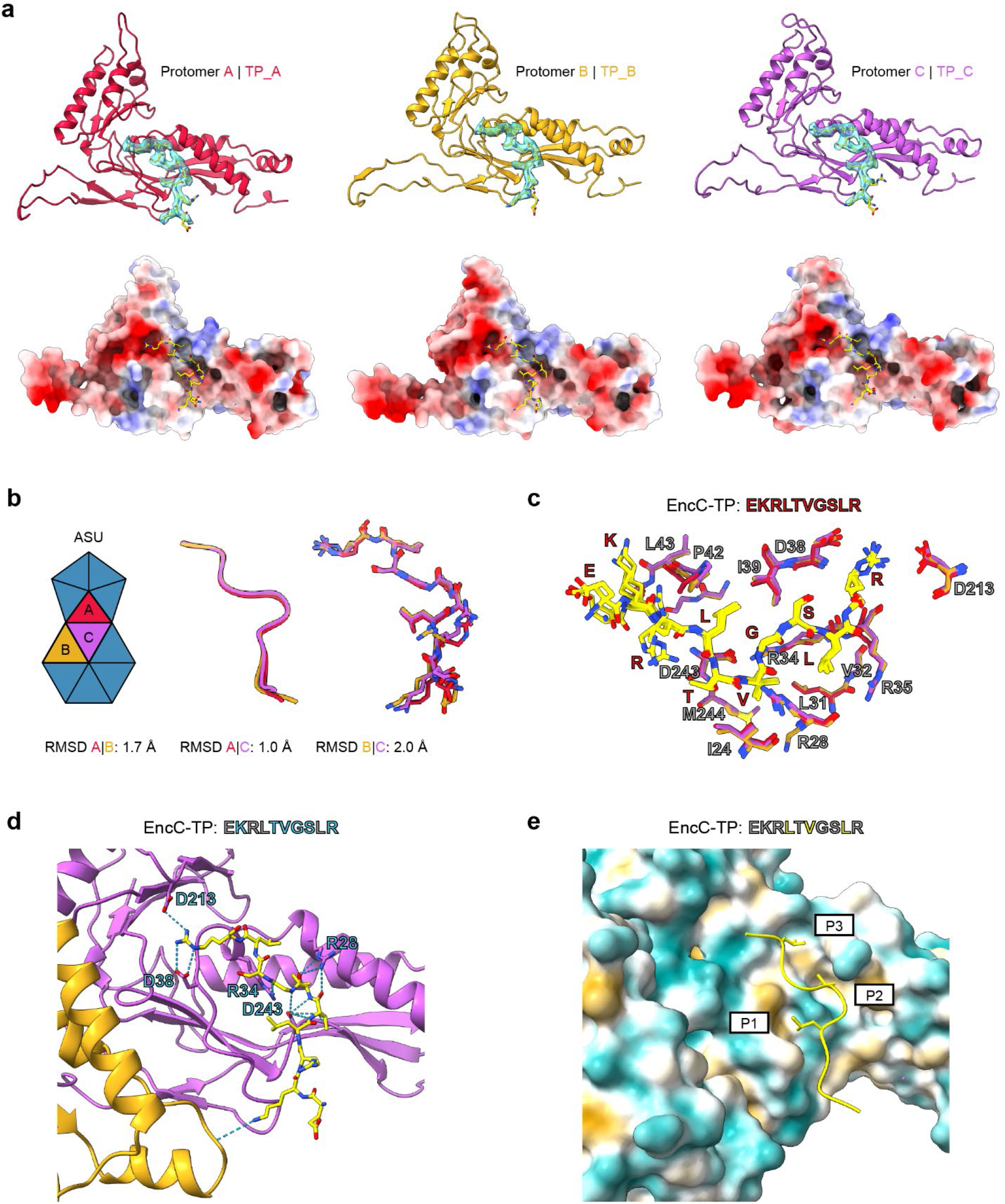
Structural analysis of TP-shell interactions in T3 encapsulins. (**a**) The interactions of the three unique TPs with their respective binding sites in the A, B, and C protomers (top) and their electrostatic surfaces (bottom) with bound TPs (stick) are shown. (**b**) ASU diagram (left) and structural alignment of TP_A, TP_B, and TP_C highlighting their highly similar conformations (middle: ribbon, right: stick). Root-mean-square deviations (RMSDs) of the TPs are shown. (**c**) Structural alignment of the A, B, and C TP binding sites with bound TPs. The conformations of the TP core sequence (RLTVGSL) are highly similar between TP_A, TP_B, and TP_C. (**d**) Representative TP-binding site (TP_C) highlighting ionic and H-bonding interactions. (**e**) The same view as in **d** highlighting hydrophobic TP-shell interactions. The three hydrophobic pockets P1, P2, and P3 are shown together with the corresponding hydrophobic residues mediating the interaction.

## Discussion

All characterized natively loaded encapsulins assemble into only one type of defined icosahedral shell of T1, T3, or T4 geometry (Giessen, 2022). The only encapsulin studied to date known to exhibit substantial amounts of structural polymorphism under certain conditions is the *M. xanthus* Family 1 encapsulin. The Mx encapsulin system which natively forms 32 nm T3 shells assembled from the shell protein EncA and loaded with three cargo proteins – EncB, EncC, and EncD – is known to form 18 nm T1 shells as a minor assembly product (20-30%) when heterologously expressed in *E. coli* without cargo co-expression (Eren et al., 2022; McHugh et al., 2014). When heterologously co-expressed with the native cargo proteins EncB or EncC (cargo loading: 30-40%), only cargo-loaded T3 shells are observed (Eren et al., 2022). Similarly, when co-expressed with non-native cargo proteins, exclusively T3 shells are observed (Lau et al., 2018). These observations indicate that the presence of cargo generally prevents the formation of empty T1 shells. It appears that under native and the tested heterologous co-expression conditions, TP-binding to EncA is faster than complete T1 shell assembly. The mechanistic details of how the presence of cargo encourages Mx T3 shell formation is currently unknown. However, it can be speculated that TP-binding results in stabilization of the E-loop conformation observed in T3 protomers which is markedly different than the one found in T1 shell protomers (Giessen, 2022). Apparently, only in the T3 E-loop conformation can protomers form both pentameric and hexameric facets, necessary for T3 shell formation. The incorporation of 20 hexameric facets in T3 shells – in addition to the 12 pentameric facets needed to form an icosahedral structure – appears to locally decrease shell curvature favoring T3 shell formation. Further, the presence of cargo during shell self-assembly might exert a negative steric effect preventing the assembly of small high-curvature T1 shells. Here, we find that in addition to discrete polymorphic shells, a second type of structural heterogeneity in the Mx encapsulin system is the formation of non-icosahedral distorted shells (Fig. 1e and Fig. 2). High cargo loading of non-native cargo – 95% in the case of SNAP-tag-TP, corresponding to over 170 occupied TP-binding sites out of 180 (Fig. 1c) – is likely the cause of the observed aberrant shell assemblies. Distorted shells are still composed of both pentameric and hexameric facets, however, their tiling deviates from the regular arrangement observed in icosahedral shells. This leads to a distorted appearance in 2D class averages and aberrant low-resolution densities in ab-initio volumes (Fig. 2c). Our results highlight that excessive cargo loading may not lead to optimally functional encapsulins, especially in a non-native context. These insights should be useful for future attempts at engineering encapsulin systems with optimized functionality for biomedical or biotechnological applications.

Cargo loading in encapsulin systems has recently been reviewed (Jones et al., 2023a). While the cargo loading mechanism in Family 1 encapsulins is fairly well understood, little information about how TDs mediate cargo encapsulation in Family 2 systems is currently available (Benisch et al., 2023; Nichols et al., 2021). The Mx encapsulin system has served as a model system for studying the molecular properties of Family 1 encapsulins and for encapsulin engineering in general, however, the previously published cryo-EM-based structures exhibited only limited resolution and map quality making it difficult to derive the details of the TP-shell interaction (Eren et al., 2022). Based on our high cargo loading system, we obtained a higher quality cryo-EM map of the TP-EncA interaction in the context of the Mx T3 icosahedral shell. We specifically focused on the TP of the main cargo EncC. We highlight a mixed ionic/H-bonding and hydrophobic mode of TP-binding (Fig. 5d,e). TP-shell interaction has previously been suggested to primarily rely on hydrophobic interactions (Eren et al., 2022; Jones et al., 2023a). Our structure shows that binding of three hydrophobic residues (EKR**L**_**(1)**_T**V**GS**L**_**(2)**_R) to three distinct hydrophobic pockets (P1, P2, and P3) is clearly important in mediating binding (Fig. 5e). Further, we highlight for the first time that an ionic interaction with a neighboring shell protomer likely contributes to the overall TP-shell interaction (Fig. 5d). Taken together, the results presented should be helpful in future attempts at modulating TP-shell interaction through rational engineering approaches.

## Abbreviations

HK97: Hong Kong 97
TD: targeting domain
TP: targeting peptide
Mx: Myxococcus xanthus
TEM: transmission electron microscopy
Cryo-EM: cryo-electron microscopy
ASU: asymmetric unit
RMSD: root mean square deviation

## CRediT authorship contribution statement

**SK:** Conceptualization, data curation, formal analysis, methodology; **MPA:** Data curation, formal analysis, methodology; **TWG:** Conceptualization, data curation, formal analysis, funding acquisition, methodology, project administration, resources, supervision, visualization, writing – original draft, review & editing.

## Data availability

Data will be made available on request.

## Acknowledgements

We gratefully acknowledge funding from the National Institutes of Health (R35GM133325). Research reported in this publication was supported by the University of Michigan Cryo-EM Facility (U-M Cryo-EM). U-M Cryo-EM is grateful for support from the U-M Life Sciences Institute and the U-M Biosciences Initiative. Molecular graphics and analyses were performed with UCSF ChimeraX, developed by the Resource for Biocomputing, Visualization, and Informatics at the University of California, San Francisco, with support from National Institutes of Health R01GM129325 and the Office of Cyber Infrastructure and Computational Biology, National Institute of Allergy and Infectious Diseases.

## Appendix A. Supplementary data

Supplementary data to this article can be found online.

