## Supplementary Material for "Structure and heterogeneity of a highly cargo-loaded encapsulin shell"

### Table of Contents

Table S1: DNA and protein sequences of EncA and SNAP-tag-TP……………………………….2

Figure S1: Cryo-EM workflow and local resolution estimation……………………………………3

**Table S1. DNA and protein sequences of the constructs/proteins used in this study.** Start and stop codons are underlined. Linker sequences are shown in red. The targeting peptide (TP) sequence is shown in cyan.

| **Name** | **DNA sequence** |
|---|---|
| **EncA** | ATGCCGGACTTTCTGGGGCATGCGGAAAACCCACTCCGGGAGGAAGAGT<br>GGGCTCGCCTTAACGAGACAGTTATTCAGGTTGCCCGTCGTTCGCTTGT<br>GGGGCGTCGGATTCTTGATATTTATGGCCCTTTGGGCGCAGGCGTCCAA<br>ACAGTACCATATGACGAATTTCAGGGTGTGAGCCCAGGCGCAGTAGACA<br>TCGTCGGGGAACAAGAAACTGCTATGGTCTTCACCGACGCCCGGAAGTT<br>CAAAACTATCCCTATCATTTACAAAGACTTCCTCCTTCATTGGCGTGACAT<br>CGAAGCTGCGCGCACGCATAATATGCCTCTTGATGTAAGTGCGGCCGCC<br>GGTGCAGCTGCCCTTTGCGCCCAGCAAGAGGATGAGCTTATCTTCTATG<br>GCGACGCTCGGCTCGGGTATGAAGGCCTTATGACGGCGAACGGTCGGC<br>TGACTGTTCCATTAGGTGACTGGACTTCCCCGGGTGGCGGCTTCCAGGC<br>CATTGTCGAAGCCACTCGTAAGTTAAACGAACAAGGCCACTTTGGTCCAT<br>ACGCTGTTGTGCTGTCACCTCGCTTATATTCCCAGTTACATCGGATTTAC<br>GAAAAAACAGGGGTCTTAGAGATCGAGACAATTCGCCAGCTCGCCTCAG<br>ATGGTGTTTATCAGTCGAATCGGTTACGGGGTGAGAGTGGCGTCGTGGT<br>CTCTACAGGCCGTGAAAACATGGATTTAGCGGTGAGTATGGATATGGTTG<br>CAGCCTACTTAGGGGCATCCCGGATGAATCACCCTTTTCGCGTACTGGAA<br>GCCCTCCTTTTGCGCATCAAGCATCCTGACGCGATCTGTACGTTAGAAGG<br>TGCTGGTGCGACTGAGCGGCGCTGA |
| | **Protein sequence** |
| **EncA** | MPDFLGHAENPLREEEWARLNETVIQVARRSLVGRRILDIYGPLGAGVQTVP<br>YDEFQGVSPGAVDIVGEQETAMVFTDARKFKTIPIIYKDFLLHWRDIEAARTH<br>NMPLDVSAAAGAAALCAQQEDELIFYGDARLGYEGLMTANGRLTVPLGDWT<br>SPGGGFQAIVEATRKLNEQGHFGPYAVVLSPRLYSQLHRIYEKTGVLEIETIR<br>QLASDGVYQSNRLRGESGVVVSTGRENMDLAVSMDMVAAYLGASRMNHPF<br>RVLEALLLRIKHPDAICTLEGAGATERR |
| | **DNA sequence** |
| **SNAP-tag-TP** | ATGGGTCCAGGTAGTGATAAAGATTGCGAAATGAAACGCACTACCTTAGA<br>TTCCCCGCTCGGTAAGTTGGAGTTGAGCGGGTGCGAACAAGGCTTGCAT<br>GAAATCATTTTTCTGGGTAAGGGCACGTCAGCAGCAGATGCAGTAGAAGT<br>GCCAGCCCCGGCGGCTGTTCTGGGGGGCCCAGAGCCGCTGATGCAAGC<br>TACCGCATGGCTCAATGCCTATTTTCACCAGCCTGAGGCTATCGAGGAGT<br>TTCCTGTGCCTGCGCTTCACCATCCGGTCTTCCAGCAGGAGTCGTTTACT<br>CGGCAGGTACTGTGGAAGCTGTTGAAGGTTGTCAAGTTTGGGGAAGTAA<br>TCTCATATAGTCACTTAGCCGCGTTAGCCGGTAATCCTGCGGCTACAGCC<br>GCGGTGAAGACTGCTTTAAGCGGGAACCCAGTTCCTATCCTTATCCCATG<br>CCACCGCGTGGTACAAGGCGACCTCGACGTGGGCGGGTATGAGGGTGG<br>GTTGGCGGTCAAGGAATGGTTGCTGGCCCATGAAGGGCATCGGTTGGG<br>GAAACGTGGTGGGGGTAGTGGCGGGGGTAGCCCAGAAAAGCGCTTGAC<br>AGTAGGGTCTCTCCGGCGTTGA |
| | **Protein sequence** |
| **SNAP-tag-TP** | MGPGSDKDCEMKRTTLDSPLGKLELSGCEQGLHEIIFLGKGTSAADAVEVPA<br>PAAVLGGPEPLMQATAWLNAYFHQPEAIEEFPVPALHHPVFQQESFTRQVL<br>WKLLKVVKFGEVISYSHLAALAGNPAATAAVKTALSGNPVPILIPCHRVVQGD<br>LDVGGYEGGLAVKEWLLAHEGHRLGKRGGGSGGGSPEKRLTVGSLRR |

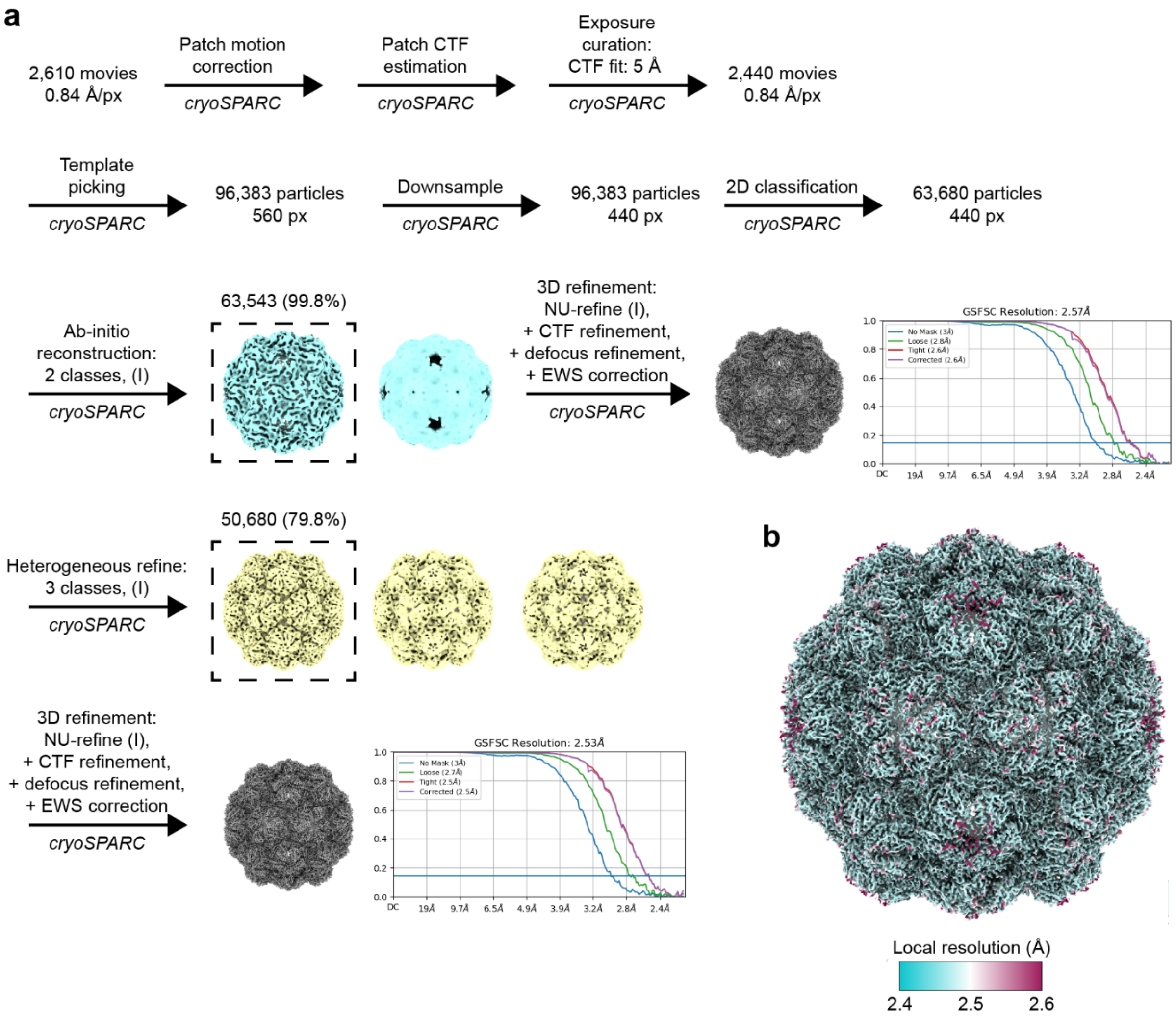


**Figure S1. Cryo-EM workflow and local resolution estimation.** (**a**) Cryo-EM workflow for the SNAP-tag-TP loaded Mx T3 encapsulin shell. (**b**) Exterior view of the Mx T3 shell colored by estimated local resolution.
